# Effects of structural complexity and water depth on the juvenile blue crab *Callinectes Sapidus* in a simulated salt marsh mesocosm

**DOI:** 10.1101/2023.01.23.524977

**Authors:** A. Challen Hyman, Cole R. Miller, Daniel Shi, Romuald N. Lipcius

## Abstract

The blue crab (*Callinectes sapidus*) is ecologically and economically important in Chesapeake Bay. Nursery habitats, which disproportionately contribute individuals to the adult segment of populations, are essential to blue crab population dynamics. *Spartina alterniflora* salt marshes are productive but fragmented intertidal nursery habitats which may serve as a refuge from predation for juvenile blue crabs. However, the effects of various characteristics of salt marshes on nursery metrics, such as survival, have not been quantified. Using mesocosm experiments, we examined the effects of shoot density and water depth on juvenile blue crab survival using adult blue crabs as predators. Survival increased significantly with shoot density, whereas water depth did not affect survival. Thus, in contrast to several previous studies, water depth did not influence survival of juvenile blue crabs, possibly due to different environmental conditions from prior studies. These findings indicate that salt marsh structural complexity enhances juvenile survival, and that the beneficial effect of shallow water on juvenile survival differs by environmental conditions.

## Introduction

Survival is a key determinant of population structure and dynamics. Early development stages (i.e., larvae, postlarvae, and young juveniles) typically exhibit the lowest survival rates throughout ontogeny of marine fish and invertebrates (Gosselin and Qian, 1997). Predation pressure is generally considered the dominant driver of juvenile mortality (Hoey and McCormick, 2004; McCormick and Hoey, 2004; Hixon and Jones, 2005; Almany and Webster, 2006). Predation on early life stages can therefore be a major bottleneck by limiting contributions to the adult segment of a population. Hence, estimating early life-stage survival associated with predation as well as quantifying effects of environmental variables on early life-stage survival rates are important both ecologically and for management.

Estuarine research on juvenile survival has largely centered on the refuge role of structurally complex, biogenic habitats such as submerged aquatic vegetation (e.g. seagrasses; Heck Jr, Hays and Orth, 2003; Bromilow and Lipcius, 2017), emergent vegetation (e.g. salt marshes and mangrove swamps; Minello et al., 2003; Sheridan and Hays, 2003), or reef-forming foundation species (e.g. coral reefs; Dahlgren et al., 2006). Juvenile survival is positively related to habitat quality and quantity because it reduces predation efficiency (Beck et al., 2001; Heck Jr, Hays and Orth, 2003; Minello et al., 2003; Adams et al., 2006). Specifically, physical habitat characteristics, such as structural complexity, modify a habitat’s refuge capacity by altering encounter or capture rates of predators with prey (Lipcius and Hines, 1986; Eggleston et al., 1990; Lima and Dill, 1990; Seitz et al., 2001; Pirtle, Eckert and Stoner, 2012). However, not all structure provides equally beneficial refuge. Variation in habitat characteristics may promote or inhibit juvenile survival both within specific habitats (e.g. salt marsh; Minello, Rozas and Baker, 2012) as well as among habitats (e.g. among salt marshes, seagrass beds, and oyster reefs; Lipcius et al., 2005; Whitfield, 2017; zu Ermgassen et al., 2021). Within-habitat differences in shoot density (Bromilow and Lipcius, 2017) and patchiness (Hovel and Lipcius, 2001, 2002) can lead to vastly different survival rates for refuge-seeking juveniles.

Water depth may also influence survival. Studies on juvenile abundance in habitats of simple structure established that small juveniles were abundant in shallow water where larger predators were absent or sparse (Ruiz, Hines and Posey, 1993; Baker and Sheaves, 2006) due to negative associations between prey survival and water depth (Ruiz, Hines and Posey, 1993; Paterson and Whitfield, 2000; Ryer, Laurel and Stoner, 2010). These observations led to the development of the ‘Shallow Water Refuge Hypothesis’ or ‘Shallow Water Refuge Paradigm’, which posits that predation pressure is lower in shallow water relative to deeper, adjacent waters and results in a refuge for small juveniles (Ruiz, Hines and Posey, 1993; Baker and Sheaves, 2007; Ryer, Laurel and Stoner, 2010).

Two mechanisms have been proposed which engender the Shallow Water Refuge Hypothesis. First, foraging efficiency of large aquatic predators diminishes in shallow water because of physiological stress either directly related to low water levels (i.e. exposure) or indirectly (e.g. increased thermal stress and decreased dissolved oxygen relative to deeper waters; Hackney, Burbanck and Hackney, 1976). Second, larger predators are vulnerable to piscivorous shorebirds and mammals and therefore avoid shallow water habitats (Ruiz, Hines and Posey, 1993; Zanette and Clinchy, 2019). However, there have been few attempts to experimentally test these proposed mechanisms (Baker and Sheaves, 2007).

Salt marshes are important, structurally complex habitats of estuarine ecosystems in North America. Vegetative structure – dense *Spartina alterniflora* shoots and roots – along tidal marsh shorelines serves as a refuge for a broad suite of prey species and corresponds with high secondary production (Craig and Crowder, 2002; Minello et al., 2003). Access to salt marsh vegetation for aquatic organisms is controlled by marsh flooding and tidal regimes. As a result, the importance of marsh structural complexity in governing survival is likely regulated by hydrology (Minello, Rozas and Baker, 2012). In mid-Atlantic estuaries, tidal amplitudes render marsh surfaces exposed for longer intervals than microtidal systems in the Gulf of Mexico. Aquatic organisms must either leave the vegetated marsh or seek shelter in residual tide pools along the pitted marsh surface (Allen, Ogburn and Kenny, 2017). Prey survival is likely a function of both structural refuge as well as shallow water refuge, with the relative importance of each determined by tidal amplitudes, marsh surface inundation times and species characteristics (de la Barra et al., 2022).

Several ecologically and economically important species, including the blue crab *Callinectes sapidus*, utilize salt marsh habitats. Salt marshes serve as an alternative nursery habitat for juvenile blue crabs in locations where seagrass – the blue crab’s preferred nursery habitat – is absent or declining (Jivoff and Able, 2003; Bishop et al., 2010; Johnson and Eggleston, 2010). Salt marshes may also serve as high quality nurseries even where seagrasses are present, particularly in microtidal locations with prolonged inundation intervals (e.g. the Gulf of Mexico; Thomas, Zimmerman and Minello, 1990; Rozas and Minello, 1998; Heck Jr, Coen and Morgan, 2001). Similar to other estuarine-dependent species, salt marshes afford refuge to juvenile blue crabs through structurally complex shoots and rhizomes (Fitz and Wiegert, 1991; Zimmerman, Minello and Rozas, 2002; Minello et al., 2003; Johnson and Eggleston, 2010; Isdell et al., 2021) as well as through detrital material along erosional marsh shorelines (Etherington and Eggleston, 2000; Etherington, Eggleston and Stockhausen, 2003; Voigt and Eggleston, 2022). Blue crab secondary production is enhanced by availability of fringing salt marsh edge (Zimmerman, Minello and Rozas, 2002; zu Ermgassen et al., 2021; Hyman et al., 2022), especially in locations devoid of structurally complex, submersed macrophytes (Johnson and Eggleston, 2010; Johnston and Caretti, 2017). However, the degree to which salt marsh structural complexity promotes survival in juvenile blue crabs was contradictory in mesocosm experiments using artificial *S. alterniflora* shoot densities. In one, mortality of post-larvae or first-instar juveniles preyed on by *Fundulus heteroclitus* did not differ by shoot density (Orth and van Montfrans, 2002). In another, high *S. alterniflora* shoot densities enhanced survival of similarly-sized juvenile crabs cannibalized by larger conspecifics (Johnston and Caretti, 2017). These conflicting results may be due to methodological differences or the choice of predator. Moreover, the results were likely size-specific, as juvenile blue crabs reached a size refuge from *F. heteroclitus* at 12 mm carapace width (Orth and van Montfrans, 2002).

Structural complexity and water depth in intertidal salt marshes may collectively provide high quality refuge for juvenile blue crabs. In tethering studies, juvenile blue crab survival was inversely related to water depth (Ruiz, Hines and Posey, 1993; Dittel et al., 1995; Hines and Ruiz, 1995). Thus, intertidal salt marsh creeks may enhance survival across the tidal regime – in structurally complex *S. alterniflora* shoots at flood tide and in adjacent shallow subtidal waters at ebb tide.

Despite the clear effects of shallow water on juvenile blue crab survival, the underlying mechanisms driving this phenomenon remain uncertain. Juvenile blue crab survival and the proportion of damaged, surviving juvenile crabs both increase in shallow water, suggesting that predator foraging efficiency is reduced in these conditions (Hines and Ruiz, 1995). However, the primary predator of juvenile blue crabs is conspecific adults (Hines and Ruiz, 1995; Moody, 2003; Eggleston, Bell and Amavisca, 2005; Bromilow and Lipcius, 2017). Unlike piscivorous fish, adult blue crabs can withstand exposure to air for prolonged intervals without incurring adverse effects (Batterton and Cameron, 1978). Therefore, it is unlikely that shallow water alone would inhibit foraging of adult blue crabs.

In this study, we examined the effects of salt marsh structural complexity and water depth on survival for juvenile blue crabs. Under controlled laboratory conditions, we experimentally manipulated artificial salt marsh shoot density and water depth using juvenile blue crabs as prey and adult conspecifics as predators. Our objectives were to (1) ascertain the effects of increasing marsh structurally complexity on survival, (2) determine whether previously observed patterns of shallow water refuge in juvenile blue crabs are due to reductions in foraging efficiency, and (3) evaluate the extent to which structural complexity and water depth interact to promote survival.

## Logical framework

We employed mesocosms to determine the effects of water depth and salt marsh shoot density on juvenile blue crab survival. Under an Information Theoretic framework (**?**) we developed multiple alternative hypotheses (H_i_) (Chamberlain et al., 1890). Herein we describe and justify the hypotheses and corresponding independent variables.

**H_1_**: Water depth reduces survival, as predation pressure on juvenile fish and crustaceans is increased in deeper water because larger predators experience decreases in foraging success in shallow water (Ruiz, Hines and Posey, 1993; Dittel et al., 1995; Hines and Ruiz, 1995).
**H_2_**: Survival increases with marsh shoot density, based on juveniles having access to refuge from predation in structured habitats (Lipcius et al., 2001; Hill and Weissburg, 2013).
**H_3_**: Survival is a function of both water depth and shoot density. These two variables may additively promote survival (H_3a_) or may interact (H_3b_) to synergistically promote survival. All subsequent hypotheses will include an additive-only model (H_ia_) and a model with a water depth-shoot density interaction (H_ib_)
**H_4_**: Survival is a function of water depth, shoot density, and predator size, such that survival is inversely related to predator size because larger crabs may be restricted in movement more than small juveniles and as a result less efficient when foraging for small juveniles (Arnold, 1984; Blundon and Kennedy, 1982; Hill and Weissburg, 2013; Shakeri et al., 2020).
**H_5_**: Survival is a function of water depth, shoot density, predator size, and prey size. As juvenile blue crabs grow, they are less susceptible to predation as their carapace widens and becomes harder, spines become more prominent, and aggressive behavior intensifies (Hines and Ruiz, 1995; Hovel and Lipcius, 2002; Lipcius et al., 2007; Bromilow, 2017).

## Methods

### Experimental design

Eight 160-L recirculating cylindrical fiberglass tanks with a bottom area of 0.36 m^2^, aerated by an airstone, simulated an estuarine marsh. Juvenile blue crabs were caught weekly using dip-nets in local seagrass and algal habitats. Prior to the experiment, juvenile blue crabs were set into tanks without a predator and recovered 24 h later after draining the tank completely (n = 14). All animals were found within 5 min of searching, which validated the assumption that missing animals at the conclusion of a trial were eaten. In each tank, PVC aqueducts continuously supplied river-sourced water at a constant flow rate. Tank water was changed completely after each trial (i.e. every 48 h) to reduce buildup of ammonia, nitrates, and other waste compounds. Tank sand was similarly completely removed from the tank following the conclusion of a trial and replaced as an additional precaution to ensure all prey were collected. Before each trial, temperature, salinity, and dissolved oxygen (DO) were measured in each tank with a YSI data sonde to account for natural fluctuations in the river-sourced flow-through water.

Adult blue crabs were selected as model predators, as adult conspecifics are among the most important predators of small juveniles (Moody, 2003; Eggleston, Bell and Amavisca, 2005; Hines, 2007; Lipcius et al., 2007; Bromilow and Lipcius, 2017). Adult crabs were caught using crab traps in the York River. Following capture, each crab was measured, tagged, placed in a holding tank, and fed juvenile blue crabs to acclimate predators to prey. Holding tanks employed the same flow-through river water as experimental tanks. If an adult crab molted, it was allowed to harden for at least 2 d prior to the next trial. Adult crabs were acclimated to experimental conditions, in separate cages to deter antagonistic behavior, for 14 d prior to a trial. Juvenile blue crabs (prey) were acclimated in a separate tank with individual compartments to deter antagonistic behavior.

Shoot density and water depth were experimentally manipulated for each trial, while monitoring physicochemical variables. Four shoot density treatments with 0, 64, 96, and 128 (corresponding to densities of 0, 388, 582, and 776 m^−2^, respectively) were employed based on reported densities in *S. alterniflora* salt marshes (Cranford, Gordon and Jarvis, 1989; Dai and Wiegert, 1996; Chaisson, Jones and Warren, 2022). Marsh shoot densities were simulated using wooden dowels (1 cm diameter, 30.8 cm height) placed into a 40.6 cm x 40.6 cm plastic pegboard, which was buried 3-5 cm beneath sand from the York River. A control treatment included a peg board without dowels. Water depth was controlled using pre-measured PVC standpipes in the center of each tank (Fig. 1). Prior to each trial, tanks were randomly assigned to one of five inundation treatments (0, 4, 6, 12, and 24 cm) and one of four shoot density treatments. The 0 cm inundation treatment included an inundated refuge for the predator which emulated the pitted marsh surface used by adult blue crabs to forage at low tide (Johnson, 2022).

**FIG 1.**
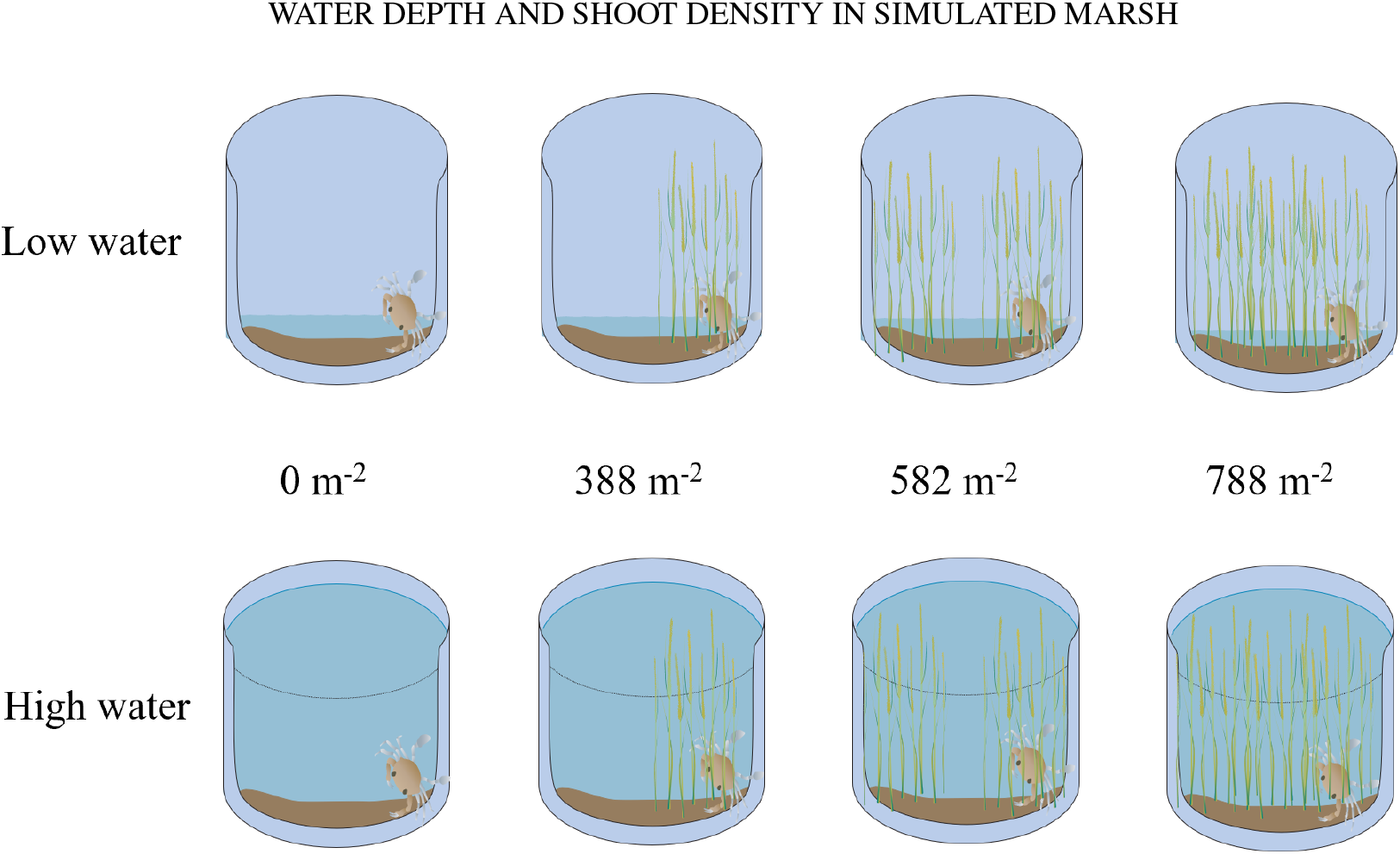
Conceptual diagram depicting mesocosm experimental design and idealized tank setup for water depth-shoot density experiment for 4 cm and 24 cm water depth treatments. Water levels for 0 cm, 6 cm, and 12 cm are not shown for brevity. Shoot density and water depth treatments were randomized regularly to avoid tank-specific bias.

For each trial, an adult blue crab (predator) was selected randomly from the holding tank and placed into each experimental tank. After recording its carapace width, a juvenile blue crab was placed in the center of each tank near the drainpipe. Each trial ran for 24 h, after which the adult crab was removed with a dip net, placed in a separate tank, fed, and left to acclimate for 24 h before the next trial. Subsequently, each tank was completely drained via a plastic siphon and searched for 10 min (i.e. twice the duration estimated to recover surviving juveniles) for surviving juveniles or carapace fragments. The absence of a juvenile crab or presence of carapace fragments was interpreted as a predation event. Then, tanks were refilled. If an adult crab died or behaved abnormally, another adult was selected randomly from a holding tank containing replacement crabs. The experiment was replicated for 17 trials (n = 136, Fig. 1).

### Data analysis

After each trial, data were recorded digitally. All data analyses, transformations, and visualizations were conducted using the R programming language for statistical computing (R Core Team, 2022). At the conclusion of each experiment, binary survival data were assessed using generalized linear regression mixed-effects models to evaluate effects of experimental treatments and water chemistry variables. The response variable, juvenile survival or death, was modeled using a binomial distribution and related to predictor variables using the logit-link (i.e. logistic regression). Predator ID, trial number, and tank ID were initially included as random-intercept effects but discarded due to negligible residual variation explained in all cases. This effectively reduced the model structures to generalized linear models.

The hypotheses for each experiment were translated into sets of statistical models (*g_i_*; Table 2) and evaluated within an information theoretic framework (Burnham and Anderson, 2002; Anderson, 2008). Salinity, temperature, and DO were initially included as fixed effects to ensure that variation in these variables did not influence survival, and were subsequently eliminated each from consideration. For each model set, AIC (Akaike’s Information Criterion) corrected for small sample size (AICc) was employed to evaluate the degree of statistical support for each model. Weighted model probabilities (w_*i*_) based on Δ_*i*_ values determined the probability that a particular model was the best-fitting model in a set. Models with Δ_*i*_ values within two points of the best fitting model were considered to have comparable support and were further evaluated using likelihood ratio *X*^2^ tests to determine their importance (Burnham and Anderson, 1998, 2002). When two models had comparable Δ_*i*_ values and likelihood ratio *X*^2^ tests did not suggest significant differences in explanatory power, the simpler model was chosen as the more appropriate model under the principle of parsimony.

## Results

In total, we ran 136 tank-trial combinations, although four trials were expunged due to outliers likely associated with erroneous equipment readings (e.g. extremely high dissolved oxygen). Data and ranges for physicochemical variables (DO, temperature, and salinity) and sizes of prey and predators are detailed in Table 1 and Figure 2.

**TABLE 1.** Summary statistics for physicochemical variables Salinity (Sal), Temperature (Temp), Dissolved Oxygen (DO), Predator Width (Predator CW), and Prey Width (Prey CW). CW = carapace width.

| Estimate | Sal | Temp<br>(°C) | DO<br>(mg/L) | Predator CW<br>(mm) | Prey CW<br>(mm) |
| --- | --- | --- | --- | --- | --- |
| Minimum | 17.08 | 23.60 | 4.68 | 104.00 | 12.40 |
| Mean | 19.83 | 26.21 | 6.09 | 132.00 | 22.32 |
| Maximum | 20.75 | 27.90 | 7.71 | 155.00 | 35.70 |

**FIG 2.**
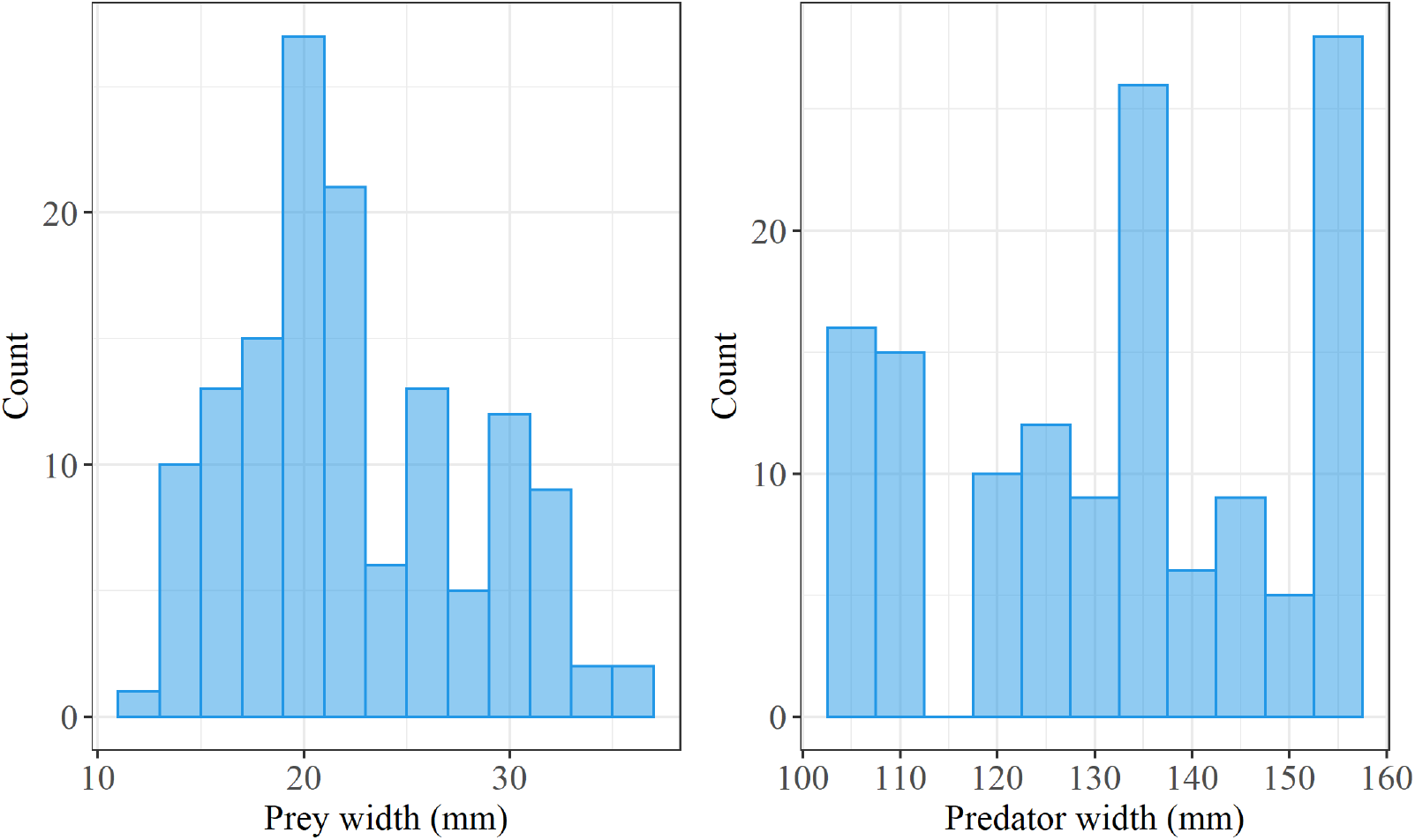
Histogram of all crab carapace widths used in the mesocosm study.

The best fitting model was *g*_2_, which posited survival solely as a function of shoot density. Models *g*_2_, *g*_3_, and *g*_5_ possessed similar AICc scores (Δ_*i*_< 2, Table 2) and a log-likelihood ratio *X*^2^ test indicated the predictive performance of these models were equivalent. However, the additional predictors in models *g*_3_ (water depth) and *g*_5_ (water depth and predator width), were not statistically significant (p > 0.05), thus we chose model *g*_2_ as the best model. The odds of survival increased positively and asymptotically with shoot density (Fig. 3) by 0.26% for every 1 unit increase in shoot density (Table 3).

**TABLE 2.**
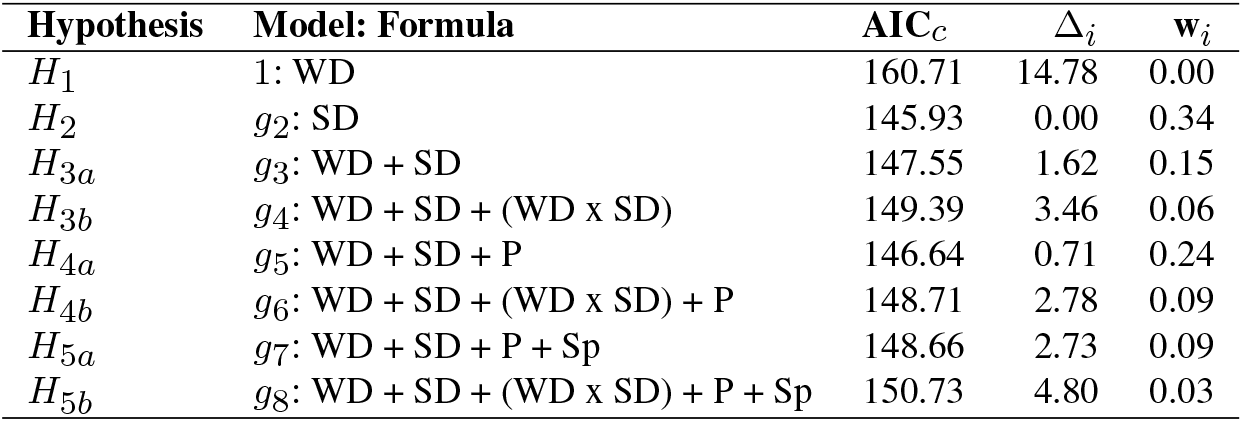
Information theoretic analysis (Anderson, 2008) of 8 logistic regression mixed-effect models (*gi*) formulated using water depth (WD), shoot density (SD), predator width (P), and prey width (Sp) as predictors of juvenile blue crab survival, where AIC*_c_* is the Akaike’s information criterion corrected for small sample size, Δ*_i_* is the difference between any model and the best model in the set, and w*_i_* is the weighted model probability that a given model is the best among the set considered.

**TABLE 3.** Summary results of model *g*_2_ coefficients: SD denotes shoot density. Estimates are on the model (i.e. logit) scale.

|  | Estimate | Std. Error | z value | P-value |
| --- | --- | --- | --- | --- |
| Intercept | -0.01 | 0.32 | -0.03 | 0.98 |
| SD | 2.6e-3 | 0.7e-3 | 3.75 | << 0.01 |

**FIG 3.**
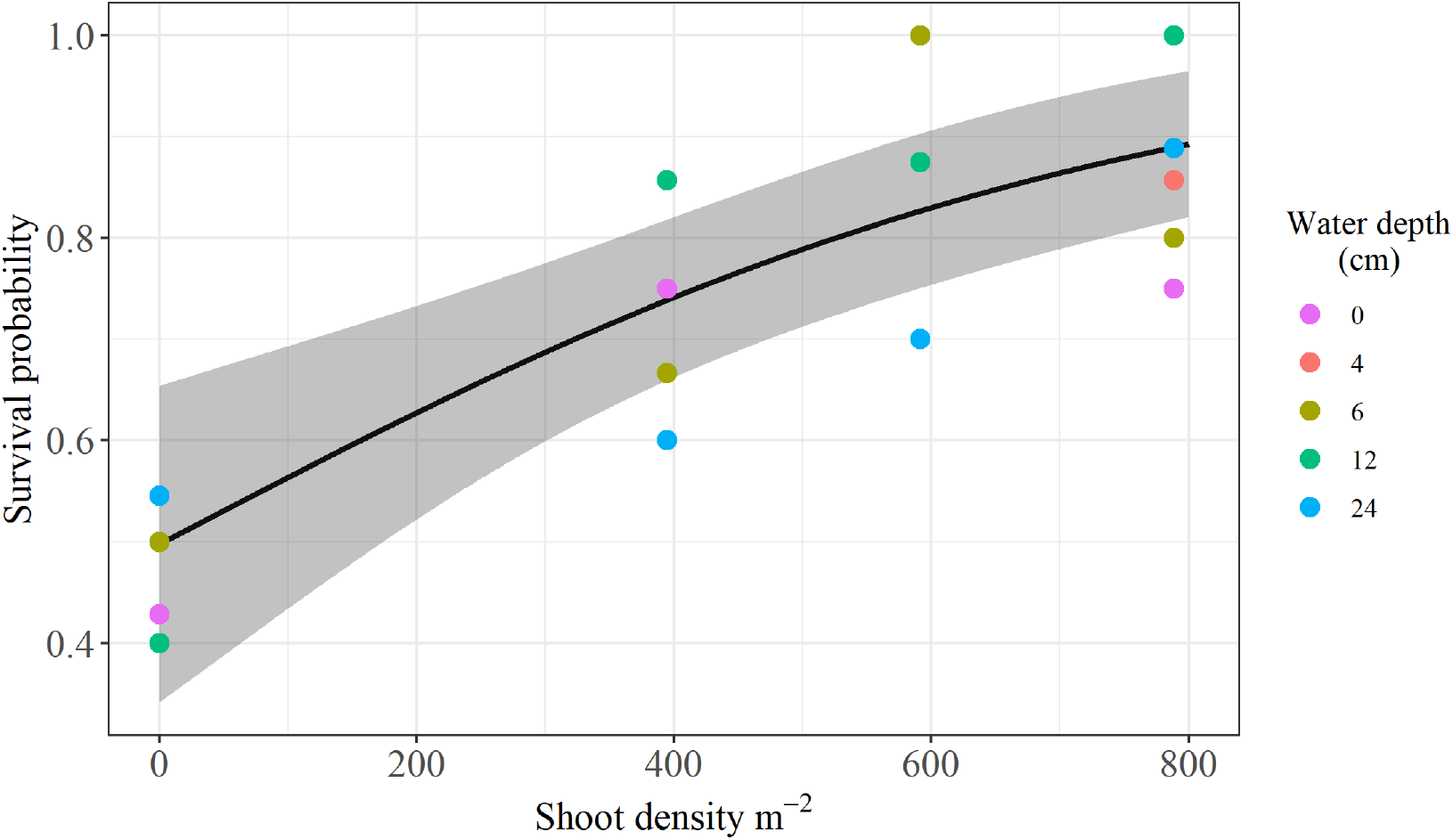
Logistic regression mean conditional effects of shoot density on survival based on estimates. Points depict aggregated data; mean survival proportions across water depth treatments and shoot densities. Shaded regions depict 95% confidence bands.

## Discussion

This study examined relationships between juvenile blue crab survival and environmental variables in simulated salt marsh habitat. Our objective was to assess the relationship between varying water depth/shoot density combinations in mediating survival of juvenile blue crabs. Multi-modal analysis indicated a relatively simple model describing survival as a function of shoot density best explained juvenile survival.

### Effects of water depth

The major findings were that survival was unaffected by water depth and that water-level effects were independent of marsh shoot density. This implies that even in areas of very shallow water, adult blue crabs experience minimal impediments to foraging. The refuge effects of shallow water may be dependent on the predator-prey interaction, such that only piscine predators too large to access shallow water are hindered (de la Barra et al., 2022). In contrast to piscine predators, adult blue crab predators are unique in their ability to withstand exposure to air, such as when adult and subadult blue crabs exploit shallow pools of muddy water at ebb tide to ambush prey when the marsh surface is exposed (Johnson, 2022).

This suggests that the second mechanism proposed to drive the shallow water refuge hypothesis – that predator behavior is inhibited by higher-level predators such as birds and mammals – underlies the observed effects of shallow water on juvenile blue crab survival with respect to cannibalizing adults (Zanette and Clinchy, 2019). For example, the presence of piscivorous shorebirds and mammals may discourage fish and adult blue crabs from entering shallow or exposed salt marsh habitat, thereby decreasing encounter probabilities and enhancing survival of small juveniles (Ruiz, Hines and Posey, 1993).

### Effects of structural complexity

Increasing shoot density enhanced juvenile blue crab survival. This effect is consistent with literature (Lipcius et al., 2001; Hovel and Lipcius, 2001, 2002; Long, Sellers and Hines, 2013) and indicates that patch-level variation in structural complexity mediates survival of juvenile crabs even at fine spatial scales and larger scales (Hovel and Lipcius, 2002). Although the effects of salt marsh shoot density on juvenile blue crab survival have been examined for smaller size classes of juveniles (e.g. 2 – 9.1 mm CW; Orth and van Montfrans, 2002; Johnston and Caretti, 2017), this is the first study describing such relationships for larger juveniles (i.e. 12 – 35 mm CW).

In salt marshes, the relationship between structural complexity and juvenile survival is likely mediated by both predator and prey size (Orth and van Montfrans, 2002; Johnston and Caretti, 2017; Hill and Weissburg, 2013). Predators optimize foraging by optimizing tradeoffs between the benefits gained and the costs incurred in obtaining prey (Hambright, 1991; Cachera et al., 2017). When prey size is too small relative to the size of the predator, the costs of pursuing the prey item relative to larger, more palatable prey items exceed the expected benefits. Hence, smaller juveniles – first-fifth instars 2.2 to 9 mm CW (Pile et al., 1996) – are most likely predated upon by smaller predators compared to larger juveniles. Small predators are less cumbered by salt marsh structural complexity because they can navigate within the interstitial spaces between shoots (Orth and van Montfrans, 2002; Hill and Weissburg, 2013). However, as juvenile outgrow the mouth-gape sizes of smaller predators, they become increasingly preferred prey to larger predators can be inhibited by salt marsh shoots and rhizomes. Although we did not detect any effect either of predator or prey size, the size ranges of both predator and prey in the present study likely reflect the latter relationship, while previous studies focusing on smaller sizes classes of both prey and predators reflect the former.

Shoot density is not the only aspect of structural complexity afforded by salt marsh habitats. Detritus exported from the vegetated marsh surface accumulates off the marsh edge and in adjacent tidal marsh creeks. This shallow detrital habitat associated with eroding peat can harbor high densities of juvenile blue crabs (Etherington and Eggleston, 2000). Shallow detrital habitats may similarly promote survival relative to unstructured sand in shallow water as well as depths accessible by piscivorous predators (Voigt and Eggleston, 2022). However, the degree to which shallow detrital habitats enhance survival in juvenile blue crabs or other prey remains untested. Future studies are required to ascertain whether the association between shallow detrital habitat and juvenile blue crabs reflects a top-down (survival) or bottom-up (growth) process (Lipcius et al., 2005; Seitz, Lipcius and Seebo, 2005).

### Relevance

Evaluating the capability of salt marshes to serve as alternative nurseries for small juvenile blue crabs has implications for blue crab population management both in the Chesapeake Bay and elsewhere. In the past, seagrass meadows were emphasized as the preferred nursery for juvenile blue crabs (Orth and van Montfrans, 1987; Perkins-Visser, Wolcott and Wolcott, 1996; Hovel and Lipcius, 2002; Ralph et al., 2013; Bromilow and Lipcius, 2017). Seagrass beds support disproportionately high densities of small (i.e. < 20mm) juvenile crabs relative to other candidate habitats (e.g. Orth and van Montfrans, 1987; Lipcius et al., 2005) and elevated survival relative to unstructured substrates (Hovel and Lipcius, 2001, 2002; Bromilow and Lipcius, 2017). However, in many locations, such as the Chesapeake Bay, seagrass meadows have been declining for decades due to anthropogenic stress Orth et al. (2010). The dominant seagrass species in the Chesapeake Bay, *Zostera marina* (eelgrass), currently experiences temperature-induced stress in summer months, giving rise to severe episodic die-offs followed by limited recovery (Moore, Shields and Parrish, 2014). These meadows are expected to continue to decline in abundance and distribution in the Chesapeake Bay as summer temperatures rise; a direct result of climate change (Waycott et al., 2009; Moore, Shields and Parrish, 2014; Wilson and Lotze, 2019). Comparisons of salt marsh nursery function to that of habitats such as seagrass meadows is necessary to understand if continued losses of *Z. marina* meadows can be compensated by salt marshes for juvenile blue crabs, as occurs in the Gulf of Mexico (Minello et al., 2003). The present study informs management in prioritizing and directing restoration and conservation efforts, as well as setting shoot density targets as metrics for restoration success.

### Caveats and future research

Inferences in this study are limited by the use of a single predator species–adult blue crabs. While the ecology of juvenile blue crabs, and specifically the importance of larger conspecifics in influencing survival, made this choice a logical first step, several piscine predators– such as blue catfish *Ictalurus furcatus*, red drum *Sciaenops ocellatus*, and striped bass *Morone saxatilis*– also consume juvenile blue crabs at high rates (Mosca III, Rudershausen and Lipcius, 1995; Hines, 2007; Lipcius et al., 2007). Hence, future studies could expand inference via replicating these experiments with a piscine predator. In relation to inundation, additional experiments could simulate tidal dynamics rather than utilizing a static system.

## Acknowledgments

The authors thank Michael Seebo for mesocosm setup and advice on capturing, acclimating, and maintaining experimental animals. Authors also acknowledge the Virginia Institute of Marine Science Research Experience for Undergraduates (VIMS REU) program. Funding for this work was provided by the National Science Foundation (grant number NSF OCE 1659656), as well as the National Marine Fisheries Service (NMFS)-Sea Grant Joint Fellowship Program in Population and Ecosystem Dynamics.

